# Recurrent *de novo* single point mutation on the gene encoding Na^+^/K^+^ pump results in epilepsy

**DOI:** 10.1101/2021.08.26.457802

**Authors:** Hong-Ming Li, Wen-Bao Hu, Chun-Gu Hong, Ran Duan, Meng-Lu Chen, Jia Cao, Zhen-Xing Wang, Chun-Yuan Chen, Fei Yin, Zhong-Hua Hu, Jia-Da Li, Li-Hong Zhong, Hui Xie, Zheng-Zhao Liu

## Abstract

The etiology of epilepsy remains undefined in two-thirds of patients. Here, we identified a *de novo* mutation of *ATP1A2* (c.2426 T>G, p.Leu809Arg), which encodes the α2 subunit of Na^+^/K^+^-ATPase, from a family with idiopathic epilepsy. This mutation caused seizures in the study patients. We generated the point mutation mouse model *Atp1a2^L809R^*, which recapitulated the epilepsy observed in the study patients. In *Atp1a2^L809R/WT^* mice, convulsions were observed and cognitive and memory function was impaired. This mutation affected the potassium binding function of the protein, disabling its ion transport ability, thereby increasing the frequency of nerve impulses. Our work revealed that *ATP1A2^L809R^* mutations cause a predisposition to epilepsy. Moreover, we first provide a point mutation mouse model for epilepsy research and drug screening.

## Introduction

Epilepsy is among the most common and widespread neurologic diseases and is characterized by recurrent seizures due to excessive discharge of cerebral neurons(Nickels et al., 2016; Vezzani et al., 2019). Epileptic patients are diagnosed with seizures together with an epileptic waveform detected using electroencephalography (EEG)(Elger & Hoppe, 2018). However, the etiological factors of epilepsy remain enigmatic. Genetic variations, metabolic disorders, cerebral tumors, and brain structure abnormalities are regarded as causative factors, and genetic variation accounts for 40% of individuals with epilepsy(Vezzani et al., 2016). However, identifying epilepsy-causative genes is difficult because the etiology of epilepsy is multifactorial and involves the influence of polygenetic variants interacting with environmental factors. Although monogenic forms of epilepsy account for 1-5% of all epilepsy cases(Striano & Minassian, 2020), identifying monogenic epilepsy genes is still challenging, since obtaining genetic and experimental evidence from both the family pedigree and animal models is not easy.

Central to genetic etiology studies, mutations of ion channels such as potassium, sodium, and calcium channels draw the most attention to unravel the underlying mechanisms of epilepsy(Aiba & Noebels, 2021; Djordjevic et al., 2021; El Ghaleb et al., 2021; Oyrer et al., 2018; Shah, 2021). These ion channels affect neuronal physiology by stabilizing and propagating neuronal activity(Oyrer et al., 2018). Genes associated with neurotransmission, neurometabolic disorders, transcriptional activation, or repression are also involved in epilepsy(Epi25 Collaborative. Electronic address & Epi, 2019). To date, thousands of genes have been reported as associated with epileptic conditions(Epi25 Collaborative. Electronic address & Epi, 2021; Fatima et al., 2021; Fry et al., 2021; Li et al., 2021; Lindy et al., 2018; Liu et al., 2018; Parrini et al., 2017; Tidball et al., 2020; Usmani et al., 2021), but most of them are not experimentally confirmed(Ran et al., 2015; Stenson et al., 2017; Takata et al., 2019; J. Wang et al., 2017). Considering that comprehensive epilepsy panels including hundreds of genes are offered by companies for genetic counseling(Poduri, 2017; Y. Wang et al., 2017), it is imperative to perform mechanistic studies and confirm genotype-phenotype associations to assist clinicians in proper diagnosis and therapeutic decision-making(Ellis et al., 2020).

*ATP1A2* encodes the α2 subunit of P-type cation transport Na^+^/K^+^-ATPase, an integral membrane protein responsible for establishing and maintaining the sodium and potassium ion electrochemical gradients across the plasma membrane(Poulsen et al., 2010). This protein pumps three sodium out of the cell and two potassium ions into the cell after the nerve impulse. This gradient, which relies on constant Na^+^/K^+^-ATPase activity, is essential for osmoregulation, sodium-coupled transport of various organic and inorganic molecules, and electrical excitability of nerve and muscle tissue (Entrez Gene: ATP1A2 ATPase). This protein is highly expressed in the brain, heart, and skeletal muscle. Structurally, the protein contains ten transmembrane helices that harbor sodium- and potassium-binding sites(Morth et al., 2007b; Shinoda et al., 2009b). Mutations in *ATP1A2* are the primary genetic cause of family hemiplegic migraine 2 (FHM2)(Du et al., 2020) and alternating hemiplegia of childhood 1 (AHC1)(Monteiro et al., 2020). Epilepsy is described as a comorbidity of FHM2 and AHC1 in *ATP1A2*- mutated cases(Costa et al., 2014; Deprez et al., 2008; Pisano et al., 2013). However, evidence to confirm whether this gene causes epilepsy is lacking. To address this question, we developed an *Atp1a2^L809R^* mouse model, which is a new genetically modified point mutation mouse model for epilepsy. The purpose of this study was to demonstrate that *ATP1A2* is an epilepsy-causative gene, providing the genetic and experimental evidence from the family pedigree to the animal model to explain the cause of epilepsy.

## Results

### Recurrent *de novo ATP1A2^L809R^* mutation in a family with idiopathic epilepsy

A family with three children were enrolled in this study. The pedigree of the family is shown in Figure 1A. Patient III-4 was born in August 2002, and was diagnosed with epileptic syndrome. Patient III-5 was born in June 2009 and died at 7 years of age with epileptic syndrome. Seizure’s onset occurred at 6 months and then twice per year on average in patient III-4 and patient III-5. The first seizures were preceded by a high fever, followed by purplish lips, glassy eyes, and dysphagia. Migraine, alternative hemiplegia, nausea, and vomiting were often associated with seizures. During the seizures, the patients lost consciousness, suffered muscular spasms, strabismus, pale skin, convulsions, drooling, dilated pupils, sweating of the palms, decreased heart rate, increased blood pressure, and bladder and bowel incontinence. Patients often slipped into comas after seizures and returned to consciousness 3 to 4 days later. Physical and mental retardation were not observed at an early age, as the patients could read, count, and recite at 2 to 3 years of age. However, mental retardation developed during aging due to repeated seizures and encephalic damage, including brain edema and atrophy (Fig. 1B). Agitation also triggered seizures. We obtained the clinical history of patient III-5. Epilepsy was examined by EEG. Low- and medium-amplitude fast spike waves were recorded in the frontal and anterior temporal areas while awake, and interictal medium spike waves were recorded in the central and occipital areas while asleep (Fig. 1C). Other members of this family did not have epilepsy. Indeed, patient III-6 was born in February 2019, without this mutation and no clinical epilepsy symptoms were observed until now.

**Fig. 1.**
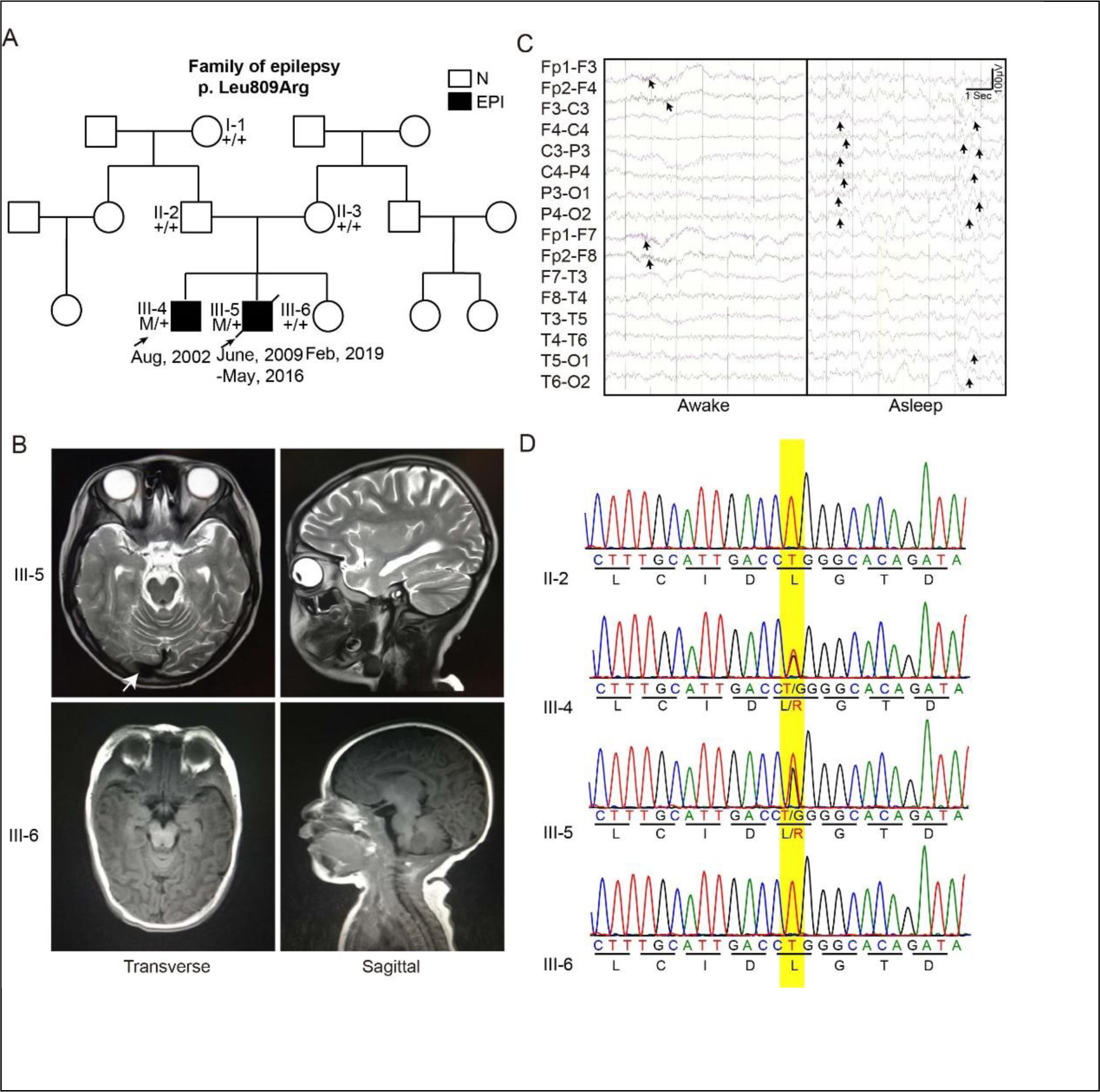
The recurrent *ATP1A2^L809R^* mutation in a family with idiopathic epilepsy. **A**, The pedigree demonstrating the *de novo* occurrence of the c.2426 T > G mutation in *ATP1A2*. I-1, II-2, II-3, III-4, III-5 were subjected to whole-exome sequencing. M, mutated allele (c.2426 T > G); +, wild-type allele; N, normal; EPI, epilepsy. **B**, MRI brain images: Abnormal signals in the white matter of the brain; the left cerebral hemisphere is slightly atrophied; the right anterior and middle cerebral arteries are altered; enlarged ventricles (7-year-old III-5). Normal MRI images (3-month-old III-6). Arrows show brain lesions. **C**, Representative electroencephalograms. Low- and medium-amplitude fast spike waves were observed in the frontal and anterior temporal lobes while awake. Medium interictal spikes were observed in the central and occipital lobes while asleep (7-year-old III-5). The patient did not cooperate with the eye opening and closing test and hyperventilation tests. Arrows show the abnormal epileptic activity in EEG recordings. **D**, Sanger sequencing demonstrating *de novo ATP1A2^L809R^* heterozygotes in III-4 and III-5. No mutation was found in III-6.

We investigated whether genetic variation was the cause of epilepsy in this family. Whole-exome sequencing was performed to screen epilepsy-causing genes (Supplementary Fig.1A). We hypothesized that the causative gene was an autosomal recessive trait that was heterozygous in their parents but homozygous in the two non-twin boys, as the two non-twin boys had epilepsy but their parents were healthy. Unfortunately, none of the annotated genes related to epilepsy met this standard (Supplementary Fig.1B) (Lindy et al., 2018; Liu et al., 2018; Parrini et al., 2017). Surprisingly, we identified a *de novo* mutation in *ATP1A2* that occurred at the same nucleotide in the two non-twin boys. We identified a c.2426 T > G variant in the *ATP1A2* gene as the only remaining candidate mutation. This missense variant changed thymine to guanine at the nucleotide position 2426 (NM_000702.4), and was predicted to result in leucine to arginine substitution (p.Leu809Arg) (NP_000693.1).

Next, we extracted DNA from blood and amplified the fragment of the *ATP1A2* gene containing c.2426T. We confirmed that the parents were normal but the two children harbored the same heterozygous c.2426 T > G mutation (Fig. 1D), which was consistent with the high-throughput sequencing results. The fact that both patients were heterozygous in the *ATP1A2* gene indicated that the L809R mutation could be a gonadal mosaicism mutation. Moreover, *ATP1A2^L809R^* is a *de novo* mutation that is absent from both the dbSNP and 1000 Genome databases, while *ATP1A2^L809R^* is reported in a patient with FHM2 in the ClinVar database without any functional evidence (accession No. VCV000529755.1). we hypothesize that *ATP1A2^L809R^* is the causative mutation for epilepsy in this family.

The third child in this family without the *ATP1A2^L809R^* point mutation is healthy, which further supports our hypothesis. We suggested the parents do amniotic fluid culture for prenatal diagnosis at 18 weeks post-conception when they had the unplanned pregnancy of the third child in 2018. Fortunately, the prenatal diagnosis from amniotic fluid culture showed that the embryo was normal without the L809R mutation (Fig. 1D). Magnetic resonance imaging (MRI) also revealed normal brain structure 3 months after birth (Fig. 1D). The third child is healthy until now (2 years of age) without any epilepsy symptoms. Follow-up studies of this child are needed.

### Recurrence of epilepsy in *Atp1a2^L809R^* point mutation mice

To determine whether the *ATP1A2^L809R^* mutation is the epilepsy-causing mutation, we generated *Atp1a2^L809R^* point mutation mice using CRISPR/Cas9 to investigate if this mouse model presented the same phenotype of the two non-twin boys (Fig. 2A). The genotype of the mice was confirmed by Sanger sequencing (Fig. 2B). To our surprise, we could only obtain mice whose genotype was *Atp1a2 ^L809R/WT^*. We observed that 8.4% of mice died after birth. All these mice were confirmed to be homozygous by Sanger sequencing (Fig. 2B, 2C), which deviated from Mendelian segregation ratios suggesting that some *Atp1a2^L809R/L809R^* mice were embryonic lethal. *Atp1a2 ^L809R/WT^* mice had seizures onset as early as 1 month after birth and died as early as 2 months (Fig. 2C, 2D; Video S1). Quantitively, 25% of *Atp1a2^L809R/WT^* mice died within 2 months. At the same time, all *Atp1a2^WT/WT^* mice were alive and healthy (Fig. 2D). Compared to the *Atp1a2^WT/WT^* mice, *Atp1a2^L809R/WT^* mice demonstrated higher frequency and amplitude epileptiform EEG activity (Fig. 2E), suggesting seizures. Overall, we found that the *Atp1a2^L809R/L809R^* point mutation was lethal *in utero* and that *Atp1a2^L809R/WT^* mice recapitulated the phenotype of *ATP1A2^L809R^* heterozygous patients.

**Fig. 2.**
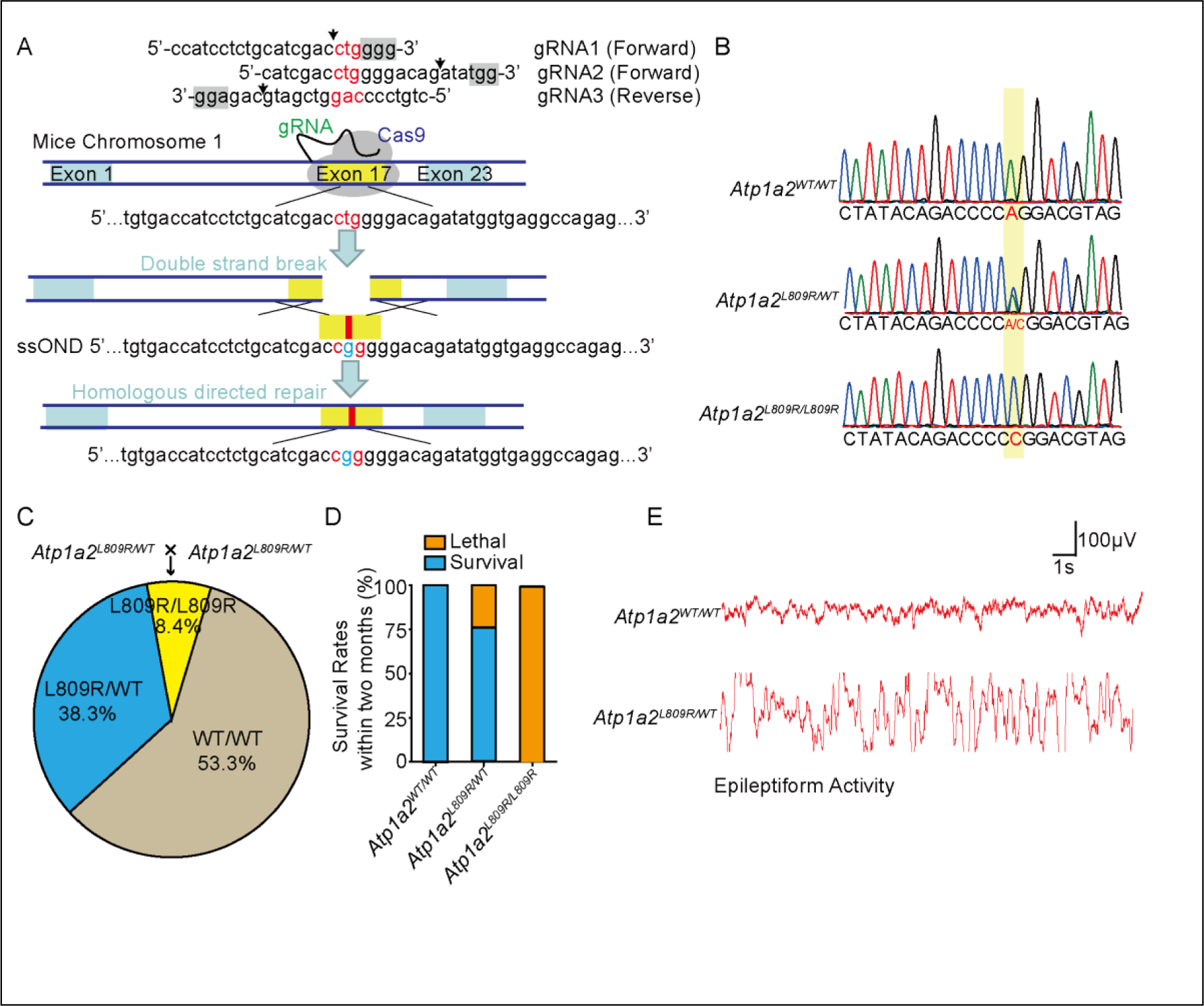
*Atp1a2^L809R^* heterozygous mice recapitulate epilepsy of the *ATP1A2^L809R^* heterozygous patients. **A**, Schematic showing the CRISPR/Cas9 techniques used to generate *Atp1a2^L809R^* point mutation mice. c.2426 T > G mutation introduced into exon 17 of mouse *Atp1a2*. Arrows showing the cutting site; grey indicates the protospacer adjacent motif (PAM) sequences; gRNA1 and gRNA2 match the forward strand DNA; gRNA3 matches the reverse strand DNA; ssOND, single-strand oligo donor DNA. **B**, Sanger sequencing demonstrating heterozygous *Atp1a2^L809R^* (*Atp1a2^L809R/WT^*) and homozygous *Atp1a2^L809R^* (*Atp1a2^L809R/L809R^*). **C**, Birth rates of *Atp1a2^WT/WT^*, *Atp1a2^L809R/WT^*, and *Atp1a2^L809R/L809R^* mice (per 50 mice). **D**, Survival rates of *Atp1a2^WT/WT^*, *Atp1a2^L809R/WT^*, and *Atp1a2^L809R/L809R^* mice within 2 months. **E**, Representative EEG demonstrating epileptiform activity in *Atp1a2^L809R/WT^* mice.

### Cognitive dysfunction and encephalic damage in *Atp1a2^L809R^* mice

To examine whether the *ATP1A2^L809R^* mutation causes dysgnosia, Morris water maze tests were performed to evaluate the cognitive function of mice. The latency to find the hidden platform was increased in the *Atp1a2^L809R/WT^* group compared with the *Atp1a2^WT/WT^* group (Fig. 3A). Further, the time spent in the target quadrant and annulus crossings through the location of the removed platform were decreased in the *Atp1a2^L809R/WT^* group compared with the *Atp1a2^WT/WT^* group (Fig. 3A). H&E staining revealed longitudinal axial atrophy in the hippocampal dentate gyrus and enlarged lateral ventricles in *Atp1a2^L809R/WT^* mice (Fig. 3B). MRI confirmed that the brain was damaged in 2-month-old mice, with enlarged lateral ventricles and small hippocampal volume in *Atp1a2^L809R/WT^* mice (Fig. 3C). Regional hypometabolism identified with ¹⁸F-FDG positron emission tomography (PET) represented the focus and projection areas of seizure activity (Duncan et al., 2016). In our case, hypometabolism was observed in the hippocampus and cerebellum of *Atp1a2^L809R/WT^* mice (Fig. 3D). Thus, *Atp1a2^L809R/WT^* mice presented cognitive decline and hippocampal damage comparable to those found in patients.

**Fig. 3.**
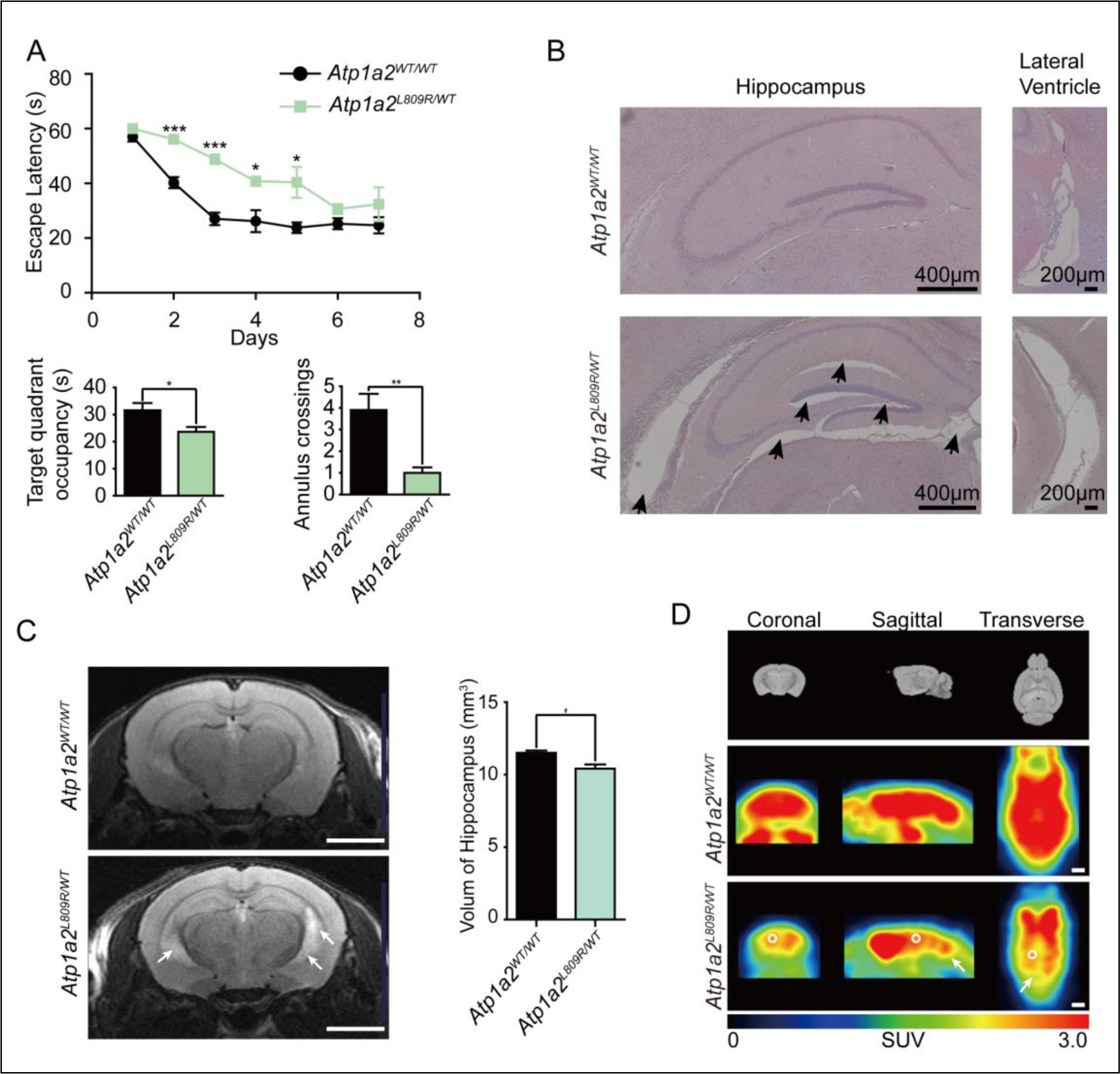
Impaired memory loss and hippocampal damage in *Atp1a2^L809R/WT^* mice. **A**, Morris water maze tests were performed to analyze the long-term memory of *Atp1a2^L809R/WT^* mice. Escape latency to the invisible platform, target quadrant occupancy, and annulus crossings were measured. *Atp1a2^L809R/WT^* mice showed a significant decrease in long-term memory performance compared to age-matched wild-type mice. n = 6. Data are presented as mean ± SD. * *P* < 0.05, *** *P* < 0.001 by two-way ANOVA with Dunnett’s post-hoc test (top); * *P* < 0.05, ** *P* < 0.01 by unpaired t-test (bottom). **B**, H&E staining showing longitudinal axial atrophy in the hippocampus and an enlarged lateral ventricle in *Atp1a2^L809R/WT^* mice. n = 6. **C**, MRI showing reduced hippocampal volume and enlarged lateral ventricles in *ATP1A2^L809R/WT^* mice. n = 5. Scale bar, 2 mm. Data are presented as mean ± SD. * *P* < 0.05 by unpaired t-test. **D**, PET-CT revealed that glucose metabolism was reduced in the hippocampus (circles) and cerebellum (arrows) of *Atp1a2^L809R/WT^* mice. n = 5. Scale bar, 2 mm.

### *Atp1a2^L809R^* mutation disrupts potassium transportation and impairs flux through the Na^+^/K^+^ pump

Next, we investigated the molecular mechanism of epilepsy caused by the *Atp1a2^L809R^* mutation. We analyzed the structure of this protein, and observed that p.Leu809 was located in the M6 transmembrane domain of ATP1A2, next to aspartic acid residues D808 and D811, which are critical for potassium-binding cavity formation (Kanai et al., 2013; Nyblom et al., 2013) (Fig. 4A, 4B). Leu 809 is located in the K^+^ binding pocket (PDB code: 4HQJ) (Nyblom et al., 2013) (Fig. 4B). The amino acid position of this variant is highly conserved in ATPase family members and is conserved across several species, including human, mouse, rat, chicken, pig, and *Xenopus* (Supplementary Figure 1C). This highly conserved region might play an important role in Na^+^/K^+^ pump function. Since leucine is a neutral amino acid, L809R mutation might add a positive charge and affect ion transport by changing the local structure of the potassium-binding cavity.

**Fig. 4.**
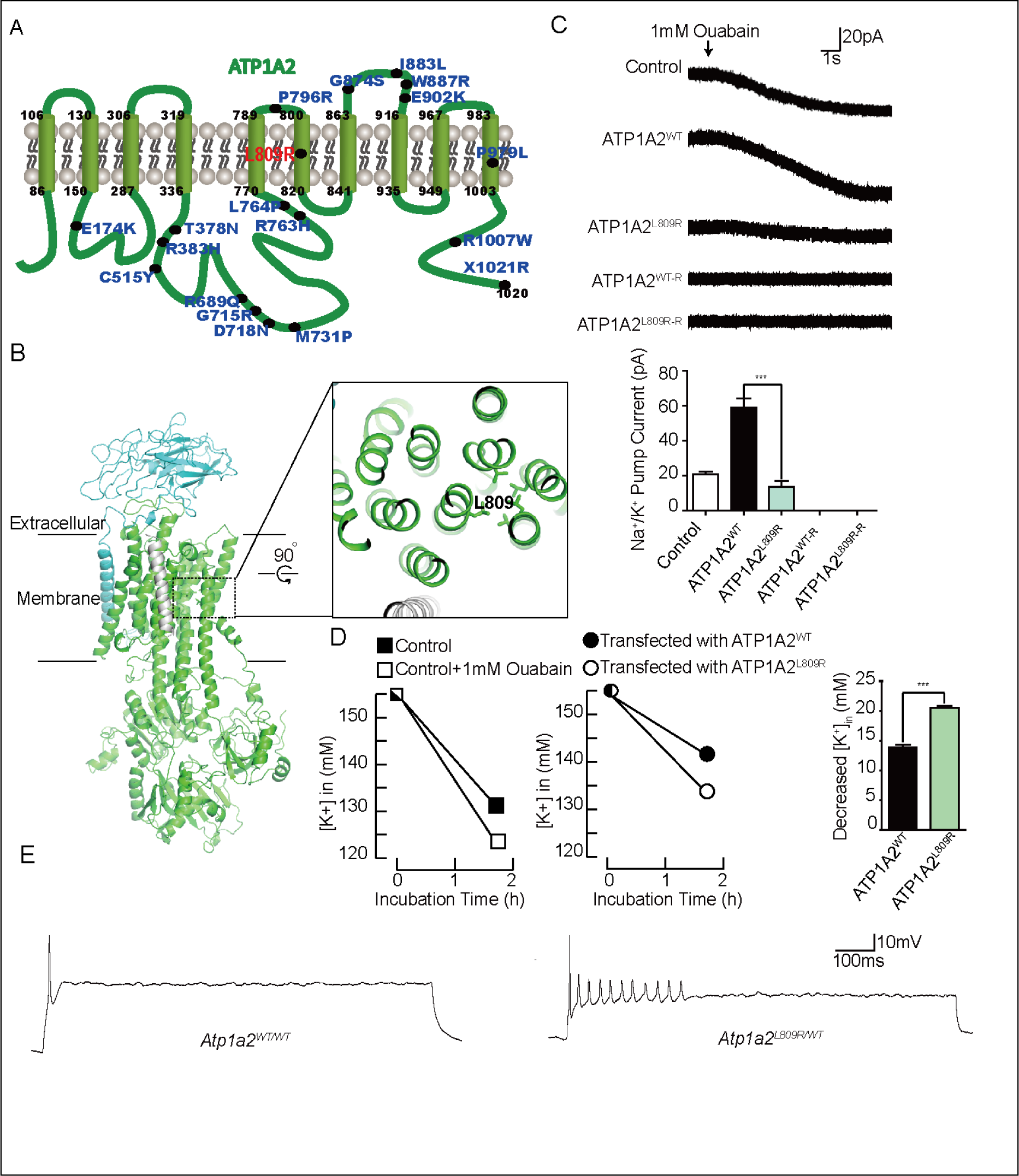
ATP1A2^L809R^ affects K^+^ transport and neuron excitability. **A**, A schematic showing the transmembrane domain structure of a single α2 subunit with ten transmembrane domains. The p.L809R substitution changes one of the leucine residues to an arginine. Other labeled sites indicate pathogenic mutations reported for ACH1 or FHM2. **B**, Leu809 is located in the K^+^ binding pocket. The structure shown here has Protein Data Bank code 4HQJ (Nyblom et al., 2013). An enlarged view showing the location of Leu 809. The positively-charged arginine substitution in the S6 segment changes the property of the transporter. **C**, Representative traces of whole-cell currents recorded in 293T cells transfected with the indicated plasmids (empty vector control, *ATP1A2^WT^*, *ATP1A2^L809R^*, *ATP1A2^WT-R^*, *ATP1A2^L809R-R^*) and treated with 1 mM ouabain. Ouabain is an inhibitor of Na^+^/K^+^ pumps, and *ATP1A2^L809R-R^* is resistant to ouabain treatment. Na^+^/K^+^ pump currents were recorded at 0 mV. n = 6. Data are represented as mean ± SD. *** *P* < 0.0001 by one-way ANOVA with Dunnett’s post-hoc test. **D**, K^+^ uptake analysis of 293T cells treated with ouabain (left) or transfected with *ATP1A2^WT^* or *ATP1A2^L809R^* (middle). Quantification of K^+^ uptake is represented (right). n = 3. **E**, ATP1A2^L809R^ causes membrane hyperexcitability in primary neurons. Representative action potential recordings of neurons isolated from *Atp1a2^WT/WT^* and *Atp1a2^L809R/WT^* mice. Current-clamp recordings were performed by injecting a suprathreshold stimulus of 200 pA for 950 ms. Representative traces are shown. n = 6 for each group.

**Fig. 5.**
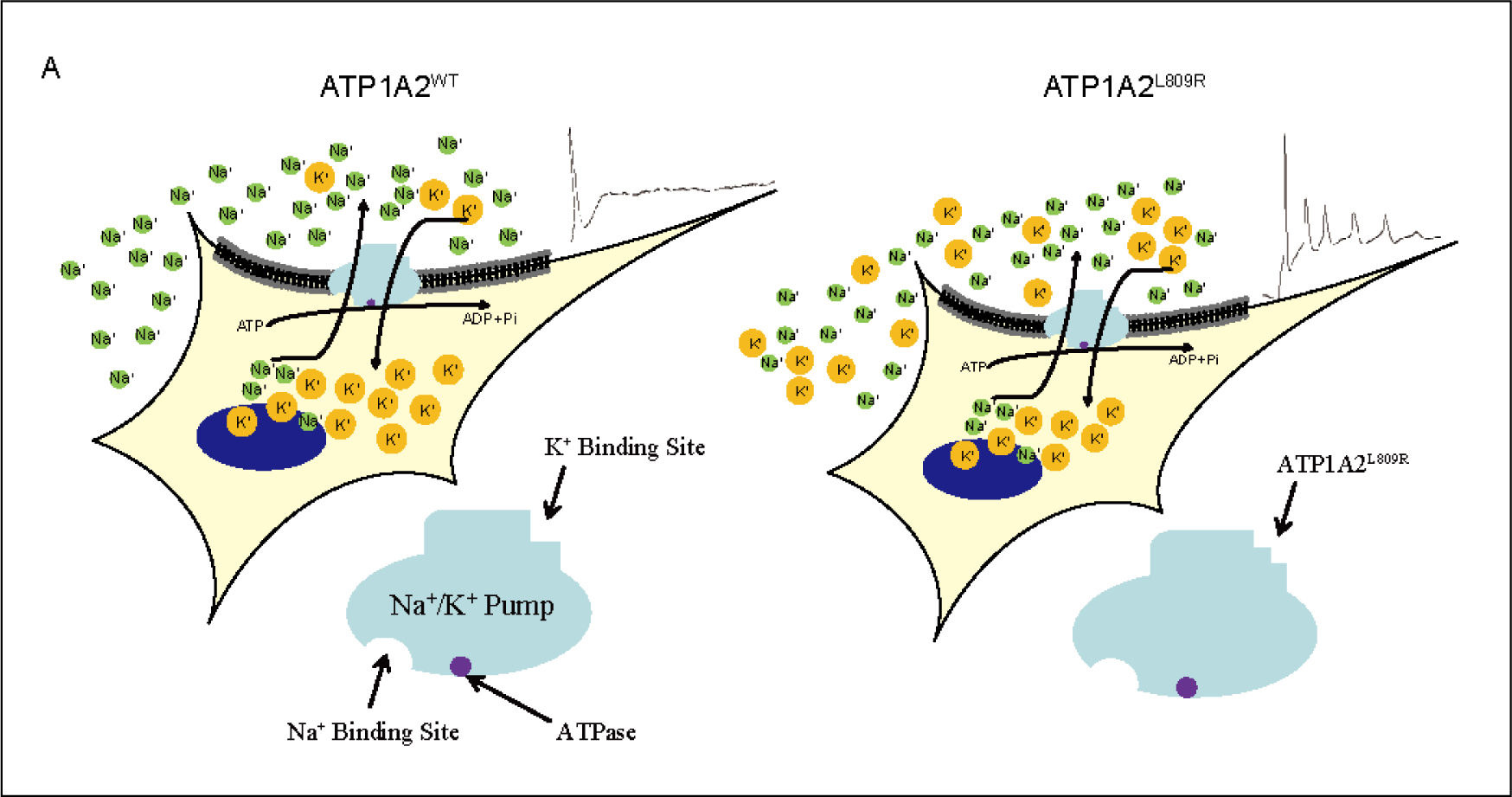
A schematic showing how ATP1A2^L809R^ affects K^+^ transport and causes epilepsy.

To answer this question, we examined Na^+^/K^+^ pump current in 293T cells overexpressing *ATP1A2^WT^, ATP1A2^L809R^*, and the ouabain-resistant mutants *ATP1A2^WT-R^* (Q116R, N127D) and *ATP1A2^L809R-R^* (L809R, Q116R, N127D). Ouabain is a Na^+^/K^+^-ATPase inhibitor. We found that the Na^+^/K^+^ pump current in the *ATP1A2^L809R^* group was lower than that in the *ATP1A2^WT^* group. The pump currents were blocked by 1 mM ouabain in the *ATP1A2^WT^* and *ATP1A2^L809R^* groups. However, the current could not be inhibited in the ouabain-resistant groups, suggesting that the blocked currents were generated by Na^+^/K^+^-ATPase and affected by L809R substitution (Fig. 4C). Protein expression and membrane distribution were not different between the *ATP1A2^WT^* and *ATP1A2^L809R^* groups, indicating that the decreased pump current in the *ATP1A2^L809R^* group did not result from altered protein expression level or membrane distribution (Supplementary Fig.2A-B). Enzyme activity was also not affected by the *ATP1A2^L809R^* mutation (Supplementary Fig.2C). These data suggest that *ATP1A2^L809R^* abrogated pump current *in vitro*. Furthermore, we measured K^+^ concentration in cells after a 2 h incubation in K^+^-free culture medium. The potassium concentration was lower in *ATP1A2^L809R^*-overexpressing cells than in *ATP1A2^WT^*-overexpressing cells (Fig. 4D). The lower K^+^ concentration inside the cells indicated aberrant potassium transport in the *ATP1A2^L809R^* mutant.

### *ATP1A2^L809R^* mutation causes neuronal hyperexcitation

The K^+^ gradient is critical for nerve impulses. Therefore, we isolated primary neurons from *Atp1a2^WT/WT^* and *Atp1a2^L809R/WT^* mice to evaluate neuronal function. We recorded action potentials and found that the action potential frequency and amplitude were enhanced in neurons from *Atp1a2^L809R/WT^* mice compared with *Atp1a2^WT/WT^* mice (Fig. 4E). These results suggest that neurons isolated from *Atp1a2^L809R/WT^* mice were hyperexcited. ATP1A2 protein expression and membrane distribution between *Atp1a2^WT/WT^* and *Atp1a2^L809R/WT^* mice were not different (Supplementary Fig.2D-E). Enzyme activity was not significantly different between *Atp1a2^WT/WT^* and *Atp1a2^L809R/WT^* mice (Supplementary Fig.2F).

### *ATP1A2* is mutated in other epilepsy cases and diseases

We identified an epilepsy case in Xiangya Hospital with a heterologous mutation in exon 18 of ATP1A2 (c.2563G>A, p.Gly855Arg). Gly855 is located within the potassium-binding pocket, implying that residues in this pocket are important for ATP1A2 function. ATP1A2^L809R^ is reported in a patient with FHM2 in the ClinVar database, without any functional evidence (accession No. VCV000529755.1). Other clinical case reports revealed epilepsy as part of the phenotype associated with ATP1A2 mutations (Monteiro et al., 2020). Moreover, frequent mutations in the ATP1A2 gene were also found in numerous human cancers according to The Cancer Genome Atlas (TCGA) datasets (Cerami et al., 2012; Gao et al., 2013) (Supplementary Fig.3). These reports suggest that *ATP1A2* is a disease-causing gene and extremely important for ion balance inside and outside cells.

## Discussion

Identifying genes that cause epilepsy and the underlying mechanisms will advance understanding of the etiology of epilepsy from a genetic viewpoint and inform therapeutic decision-making. We found that *ATP1A2* is an epilepsy-causing gene in this family with idiopathic epilepsy. Therapeutically targeting ATP1A2 may be a promising and effective translational therapy for epilepsy caused by ATP1A2 dysregulation. Mutations of the *ATP1A2* gene should be given intensive attention for preimplantation and prenatal genetic diagnosis. Further, this gene should be included in gene sets that predict epilepsy susceptibility.

Epilepsy with early-onset seizures usually leads to irreversible encephalic damage and severe physical and mental retardation. Most patients cannot take care of themselves and die at an early age due to the resulting damage. There are no effective drugs and only few surgical operations for effective epilepsy therapy(Ding et al., 2021). Therefore, prenatal diagnosis is particularly important. Screening identified epilepsy-causing genes can help avoid the birth of a child with inherited or *de novo* gene mutations that may cause epilepsy. These genes can also be explored as drug targets for epilepsy treatment. Although many epilepsy-causative *loci* have been identified(Chopra et al., 2021; Djordjevic et al., 2021; Helbig et al., 2018), the responsible genes remain to be identified.

Here, we first characterized *ATP1A2^L809R^* as an epilepsy-causing mutation through a family study and recapitulated the epilepsy features using an *Atp1a2^L809R^* point mutation mouse model. In the present study, the two non-twin boys of the family had epilepsy. The inheritance pattern of the epilepsy phenotype in this family was likely autosomal recessive inheritance. Thus, we performed whole-exome sequencing and analysis of epilepsy-related genes, but no candidate genes were observed to suggest autosomal recessive inheritance. Notably, a *de novo* mutation in *ATP1A2* in the two non-twin children drew our attention. Our case was also supported by the report from the Epi4K Consortium & Epilepsy Phenome/Genome Project, which showed a significant excess of *de novo* mutations in epileptic encephalopathies(Epi et al., 2013; Hamdan et al., 2017). Interestingly, the two non-twin boys were mutated at the same site with the same variant. One reasonable explanation was that one parent may have gonadal mosaicism. We noted that the *de novo* mutation might be inherited from parents rather than autonomously mutated in children, and that the actual risk of recurrence in families with an affected child may be as high as 50% (Myers et al., 2018). To verify this hypothesis, we extracted DNA from sperms, amplified the mutated region of *ATP1A2*, inserted the PCR products into T-vectors, sequenced 100 T-vectors and found no mutation in the sperm DNA. We could not investigate whether the mother’s oocytes were chimeric, because oocyte collection was not approved during ethical review. Therefore, we could not tell from whom the mutation was inherited. Our early study showed that *Atp1a2^L809R^* might be the epilepsy-causative mutation, so when the parents had an unplanned pregnancy, we suggested a prenatal diagnosis to exclude the L809R mutation. Fortunately, *ATP1A2* was not mutated in the third child, which was born and has been completely normal until now (2 years of age). Given that epilepsy occurred at 6 months in both patients, we have reason to believe that the child without the *ATP1A2^L809R^* mutation is healthy.

The α2 subunit of Na^+^/K^+^-ATPase consists of ten transmembrane domains and represents standard functional features of the P-type family of active cation transport proteins. The L809 residue resides in the M6 transmembrane domain, which is critical for K^+^ binding. L809R substitution appears to cause a change in ion transport. Structural analysis of the L809R variant showed that it affected potassium binding(Morth et al., 2007a; Shinoda et al., 2009a), and thus may disable the ion transport through the protein. Our data also indicated that the Na^+^/K^+^ pump current was lower in the L809R variant, which caused neuronal hyperexcitation.

Mutations in *ATP1A2* have been described in FHM2 and AHC1. These neurological disorders are dominantly inherited and are primarily caused by missense mutations(Chatron et al., 2019; Vetro et al., 2021). L764P, W887R, M731T, R689Q, D718N, P979L, E174K, C515Y, I286T, and T415M mutations occurred in families with FHM2(De Fusco et al., 2003; Jurkat-Rott et al., 2004; Todt et al., 2005; Vanmolkot et al., 2003; Vanmolkot et al., 2007). A T378N(Swoboda et al., 2004) mutation was found in patients with AHC1. Novel R1007W (Pisano et al., 2013), G874S (Costa et al., 2014), G900R, and C702T mutations were found in FHM2 with seizures. Our study expands these previous findings. Our results confirm that the L809R mutation in *ATP1A2* causes epilepsy and suggest that patients carrying *ATP1A2* mutations should be warned about the susceptibility of epilepsy.

Here, we generated a new genetically engineered mouse model of familial epilepsy. This gene-to-phenotype mouse model based on a phenotype-to-gene study enables further epileptogenic studies and antiepileptic drug screening. Epilepsy animal models need to meet three criteria: first, similar etiology to human epilepsy; second, identical physiological and genetic manifestations to human epilepsy; and third, an efficient therapeutic response to antiepileptic drugs (Grone & Baraban, 2015). Currently, there is no single animal model of epilepsy that fully represents this disease(Grone & Baraban, 2015). Researchers should match their study aims with the advantages of each model.

Acute mouse models (maximal electroshock model, MES; pentylenetetrazol model, PTZ) were used to screen antiepileptic drugs and were biased toward drugs that act on ion channels. These models appear more like seizure models rather than epilepsy models. Chronic mouse models (kindling) are used to evaluate the physiological and pathological changes in epilepsy occurrence and development, but establishing a chronic model is costly and time-consuming(Grone & Baraban, 2015). Additionally, some rat models were genetically-fixed by selective breeding, and the underlying genetic mutation responsible for epilepsy in these animals remains a mystery(Grone & Baraban, 2015). Here, we generated a new genetic epilepsy mouse model with a single point mutation. *Atp1a2^L809R^* mice showed similar etiology and pathology to those of human epilepsy, which broadens the genetically modified mouse model for epileptogenic studies and antiepileptic drug screening. Heterozygous *Atp1a2^L809R^* caused epilepsy, while the homozygous mutation was prenatally lethal similar to the findings of previous studies. Homozygous *Atp1a2* knockout mice (*Atp1a2^-/-^*) are perinatally lethal due to absent respiratory activity resulting from abnormal Cl^-^ homeostasis in brainstem neurons(Isaksen & Lykke-Hartmann, 2016).

Taken together, we identified *ATP1A2^L809R^* as an epilepsy-causing mutation in a family with idiopathic epilepsy and established an *Atp1a2^L809R^* point mutation mouse model. *ATP1A2^L809R^* should ultimately be included in data sets for prenatal diagnosis of epilepsy. Our *Atp1a2^L809R^* point mutation mouse model will have a significant impact on epileptogenic studies and antiepileptic drug screening.

## Materials and methods

### Ethics statement

All procedures involving human samples and animal care in this study were reviewed and approved by the ethical review board at the Xiangya Hospital of Central South University. All the animal experiments strictly followed the Regulations for the Administration of Affairs Concerning Experimental Animals, the Chinese national guideline for animal experiments.

### Patient family

A family exhibiting an epilepsy phenotype was identified at the Third Xiangya Hospital, Central South University, Changsha, China. A total of 5 individuals in three generations, including two affected and three unaffected members, participated in this study (Fig. 1). The probands underwent detailed clinical evaluation including medical history, video/electroencephalograph (VEGG) examination, inherited metabolic disease determination tests, and urine organic acid determination tests. All the participants were informed of the research studies and provided written informed consent.

### Sample preparation and whole-exome sequencing

Genomic DNA was extracted from peripheral blood from all the family members who participated in the study. DNA purity and concentration were measured on a Nanophotometer spectrophotometer (IMPLEN, CA, USA) (OD260/280 = 1.8∼2.0). DNA degradation and suspected RNA/protein contamination were evaluated by electrophoresis on 1% agarose gels. The concentration and purity of DNA samples were further quantified using Qubit DNA Assay Kits and a Qubit 2.0 fluorometer (Life Technologies, CA, USA).

The exome sequences were enriched from 1.0 μg genomic DNA using an Agilent liquid capture system (Agilent Sure Select Human All Exon V5). First, qualified genomic DNA was randomly fragmented to an average size of 180-280 bp using a Covaris S220 sonicator. Then, DNA fragments were end-repaired and phosphorylated, followed by A-tailing and ligation at the 3’ ends with paired-end adaptors (Illumina) with a single “T” base overhang. The DNA fragments were purified using Agencourt AMPure SPRI beads (Beckman-Coulter). The fragment size distribution and library concentrations were determined using an Agilent 2100 Bioanalyzer and qualified using real-time PCR (2 nM). Finally, the DNA library was sequenced on an Illumina Hiseq 4000 instrument (Illumina, Inc., San Diego, CA, USA) using 150 bp paired-end reads (Fig. 2).

#### Quality control

The raw image files obtained from Hiseq 4000 were processed using the Illumina pipeline for base calling and were stored in Fastq format (raw data). The reads were processed using the following quality control steps: 1) reads with adaptor contamination (>10 nucleotides aligned to the adaptor, allowing ≤ 10% mismatches) were filtered; 2) reads containing more than 10% uncertain nucleotides were discarded; and 3) paired reads with a single read having more than 50% low quality (Pared quality < 5) nucleotides were discarded. All the downstream analyses used high-quality clean data. QC statistics including total read number, raw data, raw depth, sequencing error rate, percentage of reads with average quality > Q20, percentage of reads with average quality > Q30, and the GC content distribution were calculated.

#### Read mapping

Valid sequencing data was mapped to the reference genome (UCSC hg19) using Burrows-Wheeler Aligner (BWA) software. Subsequently, Samtools and Picard ^5251585859^ were used to sort and de-duplicate the final bam file.

#### Variant calling

Reads that aligned to exon regions were collected for mutation identification and subsequent analysis. Samtools mpileup and bcftools were used for variant calling and to identify single-nucleotide polymorphisms and indels. We used CoNIFER software (Krumm et al., 2012) to identify disruptive genic CNVs in human genetic studies of disease, which might be missed by standard approaches.

#### Functional annotation

ANNOVAR software (Wang et al., 2010) was used to annotate the VCF (Variant Call Format) file obtained during variant calling. The variant position, type, conservative prediction, and other information were obtained using various databases, such as DbSNP, 1000 Genomes, ExAC, CADD (Kircher et al., 2014) and HGMD. Since we were interested in exonic variants, gene transcript annotation databases, such as Consensus CDS, RefSeq, Ensembl, and UCSC were also used for annotation to determine amino acid alterations.

#### Filtering

Variants obtained from the previous steps were filtered with MAF (minor allele frequency) > 1% using the 1000 Genomes database (1000 Genomes Project Consortium). Only SNVs occurring in exons or in canonical splice sites (splicing junction ± 10 bp) were further analyzed since we were interested in amino acid changes. Synonymous SNVs that were not relevant to amino acid alterations were discarded to get nonsynonymous SNVs that led to different gene expression products. Finally, the retained nonsynonymous SNVs were submitted to PolyPhen-2 (Adzhubei et al., 2013), SIFT, Mutation Taster, and CADD (Kircher et al., 2014) for functional prediction. SNVs identified by at least two software applications as not benign were retained.

### Analysis of potential epilepsy-causing variants

Potential candidate variants were verified by co-segregation with the phenotype within this family, based on Sanger sequencing. Primer F (introduced a 5ʹ *BamHI* site): CGGGATCCGGGGAAGAGTCCCTCTGACCTCCCTGATGCC; primer R (introduced a 3ʹ *XhoI* site): CGCTCGAGAGGGACCTGTGTGGGGTAGGAAATGGGGCAG. The PCR products (46 3bp) were analyzed using Sanger sequencing. Nucleotide sequence conservation analysis across species was analyzed using Vector NTI software.

### Generation of *ATP1A2^L809R^* point mutation mice and animal handling

The *ATP1A2^L809R^* mice were generated by Cyagen Biosciences (Guangzhou, China). We created an *ATP1A2^L809R^* point mutation in C57BL/6 mice using CRISPR/Cas9-mediated genome engineering. The mouse *Atp1a2* gene (GenBank accession number: NM_178405.3; Ensembl: ENSMUSG00000007097) is located on mouse chromosome 1. Twenty-three exons have been identified, and L809 is located on exon 17. A gRNA targeting vector (pRP[CRISPR]-hCas9-U6) and donor oligo (with targeting sequence flanked by 120-bp homologous sequences combined on both sides) were designed targeting exon 17. An L809R (CTG to CGG) mutation site in the donor oligo was introduced into exon 17 by homology-directed repair. Cas9 mRNA and gRNA generated by *in vitro* transcription and donor oligos were co-injected into fertilized eggs.

The pups were genotyped using PCR followed by sequence analysis and *HpaII* restriction analysis (wild-type allele: 730 bp and mutant allele: 419 bp and 311 bp). The gRNA target sequence and gRNA vectors were designed using VectorBuilder. gRNA1 (matches forward strand): CCATCCT(CTG)CATCGACCTG<u>GGG,</u> http://www.vectorbuilder.com/us/en/vector/VB170605-1007fsn.html; gRNA2 (matches reverse strand): CTGTCCC(CAG)GTCGATGCAG<u>AGG,</u> http://www.vectorbuilder.com/us/en/vector/VB170605-1008ekx.html; gRNA3 (matches forward strand): CATCGAC(CTG)GGGACAGATA<u>TGG,</u> http://www.vectorbuilder.com/us/en/vector/VB170605-1036add.html. Donor oligo sequence: CTGTTCATCATTGCCAACATCCCCCTTCCACTGGGCACTGTGACCATCCTCTGCATCGAC(<u>CGG)</u>GGGACAGATATGGTGAGGCCAGAGGGCAGAGTCGAGCAGTCCCACAGAATAGGGGTGGGG

### Assay of CRISPR-induced mutations

The target region of the mouse *Atp1a2* locus was amplified using PCR with specific primers: *Atp1a2*-F: TCCGATGTCTCTAAGCAGGCGG; *Atp1a2*-R:GAAGGTATCTGAAGAGGTCTGTGAAACTG. The PCR product size was 730bp; *HpaII* restriction analysis gives 730 bp or 419 bp + 311 bp from wild-type or mutant alleles, respectively. PCR products were sequenced to confirm targeting.

### Mouse electroencephalograph (EEG) recording

EEG recordings were performed to record the electrical potentials at the surface of the brain. Briefly, the mouse was anesthetized with 20 μL/g 3% pentobarbital sodium. Screws, which served as electrodes and were linked to wires, were inserted into holes drilled in the skull. The apparatus was fixed with dental cement and connected to a Multichannel Physiological Signal Acquisition and Processing System [RM-6240E] (Chengdu, China). One reference electrode was placed over the cerebellum, while the anode and cathode were implanted at (coordinates from bregma): posterior 2.3 mm, lateral 2.0 mm (left and right) separately, subdural 0.5 mm. The implants remained affixed and produced good recordings for over a month.

### Vector construction

The *ATP1A2* human cDNA sequence was cloned to the pEGFP-C1 vector between the *EcoRI* and *BamHI* restriction sites, with an additional C behind *EcoRI* to prevent frameshifts. Site-directed mutagenesis was carried out to generate *ATP1A2^L809R^* by mutating the c.2426 T>G. The ouabain-resistant form WT-R (Q116R and N127D) or triple mutants L809R-R (L809R, Q116R, N127D), were also obtained by site-directed mutagenesis.

### Patch-clamp electrophysiology

Na^+^/K^+^ pump currents (I_p_) were recorded in HEK293T cells transfected with ATP1A2^WT^, *ATP1A2^L809R^*, *ATP1A2^WT-R^*, or *ATP1A2^L809R-R^*, and treated with 1 mM ouabain. Briefly, cells were incubated in buffer (144 mM NaCl, 5.4 mM KCl, 1.8 mM CaCl2, 2 mM BaCl2, 5 mM NiCl2, 10 mM HEPES, pH 7.4 (adjusted with NaOH)) supplemented with 10 μM ouabain to block endogenous Na^+^/K^+^-ATPase. The cell membrane was patched with a pipette with inner buffer (110 mM CsCl, 40 mM NaCl, 10 mM NaOH, 3 mM MgCl2, 6 mM EGTA, 10 mM HEPES, 10 mM Mg-ATP, pH 7.4 (adjusted with CsOH)). The osmolarity was adjusted to 315 mOsm with glucose. The voltage was held at 0 mV while recording.

### Primary neuron isolation

Primary neurons were obtained from *Atp1a2^WT/WT^* and *Atp1a2^L809R/WT^* mice. Pregnant C57BL/6 mice were euthanized with 20 μL/g 3% pentobarbital sodium and E16.5 embryos were isolated. Neurons were dissected from the embryos and enzymatically digested with 2 mg/mL papain at 37 ℃ for 40 min, followed by mechanical dissociation. Isolated neurons were plated onto 100 mg/mL poly-D-lysine-coated coverslips and cultured in DMEM/F-12 medium supplemented with B27 (0.04%) and GlutaMAX (2 mM). Two days later, cells were subjected to electrophysiology recording.

### Electrophysiology recording of excitability in primary neurons

Neurons were incubated in bath buffer (140 mM NaCl, 3 mM KCl, 2 mM CaCl2, 2 mM MgCl2, 10 mM HEPES, pH 7.4 (adjusted with NaOH)). Neurons were patched with pipettes with inner buffer (140 mM KCl, 0.5 mM EGTA, 5 mM HEPES, 3 mM Mg-ATP, pH 7.4 (adjusted with KOH)). The osmolarity was adjusted to 315 mOsm with glucose.

### Potassium concentration measurements

The K^+^ concentration was measured using Cell Potassium Ion Assay Kits (cat: GMS50605.1, GenMed, USA). Briefly, HEK293T cells (1 × 10^6^ cells) overexpressing *ATP1A2^WT^* or *ATP1A2^L809R^* were incubated with K^+^-free saline in 15 mL tubes. After 2 h incubation, cells were centrifuged at 300 g for 5 min, washed three times with normal saline, and lysed with lysis buffer. The lysates were centrifuged at 16,000 g for 5 min at 4 ℃ and the protein concentration was measured. 100 μL cell lysate (200 μg/mL) was mixed with 100 μL potassium-binding benzofuran isophthalate (PFBI) and incubated in the dark for 30 min. The relative fluorescence was measured and converted to potassium ion concentration.

### Western blotting

293T cells or brain tissue extracts were incubated in RIPA buffer with proteasome inhibitors for 30 min and then centrifuged at 20,000 × g at 4 °C for 15 min to remove cellular debris. 20 µg total protein was separated using SDS-PAGE. ATP1A2 was immunoblotted with an anti-ATP1A2 (Abcam, ab166888; 1:2000).

### Morris water maze test

Morris water maze tests were performed as previously described with minor modifications. A circular pool (diameter: 120 cm and height: 50 cm) was filled with 22-23 ℃ water. The pool was divided into four quadrants of equal area. A transparent platform (diameter: 8 cm and height: 20 cm) was centered in one of the four quadrants. Four prominent visual cues were presented on each side of the pool. For visible platform tests (1.5 cm above the water surface), the water in the pool was un-dyed. For invisible platform test (1.0 cm below the water surface), the water was dyed white with non-toxic paint. Test trials were conducted for 7 days. For each daily trial, the mouse was randomly placed into the water maze at one of four quadrants. The trial was stopped when the mouse found and climbed onto the platform, and the escape latency was recorded. Visible platform tests were conducted for 4 days. A probe trial was conducted 24 h after the last acquisition session to assess the spatial retention of the location of the hidden platform. During this trial, the platform was removed from the maze, and each mouse was allowed to search the pool for 60 s before being removed.

The time spent in the target quadrant was used as a measure of consolidated spatial memory.

### Statistics

Data were analyzed using Prism 7 (GraphPad Software, San Diego, CA, USA, http://www.graphpad.com). Data are presented as mean ± standard deviation. Comparisons between groups were analyzed by unpaired t-test or ANOVA with Dunnett’s post-hoc test. *P* < 0.05 indicated statistically significant differences.

## Acknowledgements

We thank S.D.Z. at the National Institute of Biological Science, Beijing, and Z.X.L. at Sun Yat-sen University for structural analysis; W.Z. and C.N.X. at Central South University for electrophysiological recording. This work was supported by grants from the National Natural Science Foundation of China (82172502, 81974127, 81871822, 81801395), Guangdong Basic and Applied Basic Research Foundation (2016A030306051), Innovation-Driven Project of Central South University (2018CX029, 2019CX014), Science and Technology Plan Project of Hunan Province (2018RS3029), Non-profit Central Research Institute Fund of Chinese Academy of Medical Sciences (2019-RC-HL-024), and Fundamental Research Funds for the Central Universities of Central South University (2020zzts859).

## Author contributions

Z.Z.L. conceived the study, designed the experimental procedures, analyzed data, prepared the manuscript, and supervised the project. H.M.L. performed the majority of the experiments, analyzed data, and prepared the draft manuscript. W.B.H. reviewed and edited the manuscript. C.G.H, M.L.C., and R.D. performed the experiments. F.Y., and L.H.Z. contributed to clinical data acquisition. J.C., Z.X.W., and C.Y.C. participated in experimental design and provided suggestions for this project. Z.H.H., and J.D.L. helped isolating neurons and performed the electrophysiologic test. H.X. conceived the experiments, critically reviewed the manuscript and supervised the project.

## Data availability

Further information and requests for resources and reagents should be directed to and will be fulfilled by the Lead Contact, Zheng-Zhao Liu. We are glad to share the ATP1A2^L809R^ point mutation mouse generated in this study with reasonable compensation by the requestor for processing and shipping with a completed Materials Transfer Agreement. Original data of whole-exosome sequencing is available [Dryad, Dataset, https://doi.org/10.5061/dryad.tmpg4f500].

## Code availability

The patient-related data is from the clinician. All data are available from the corresponding author upon reasonable request.

## Supplementary information

**Supplementary Fig. 1.**
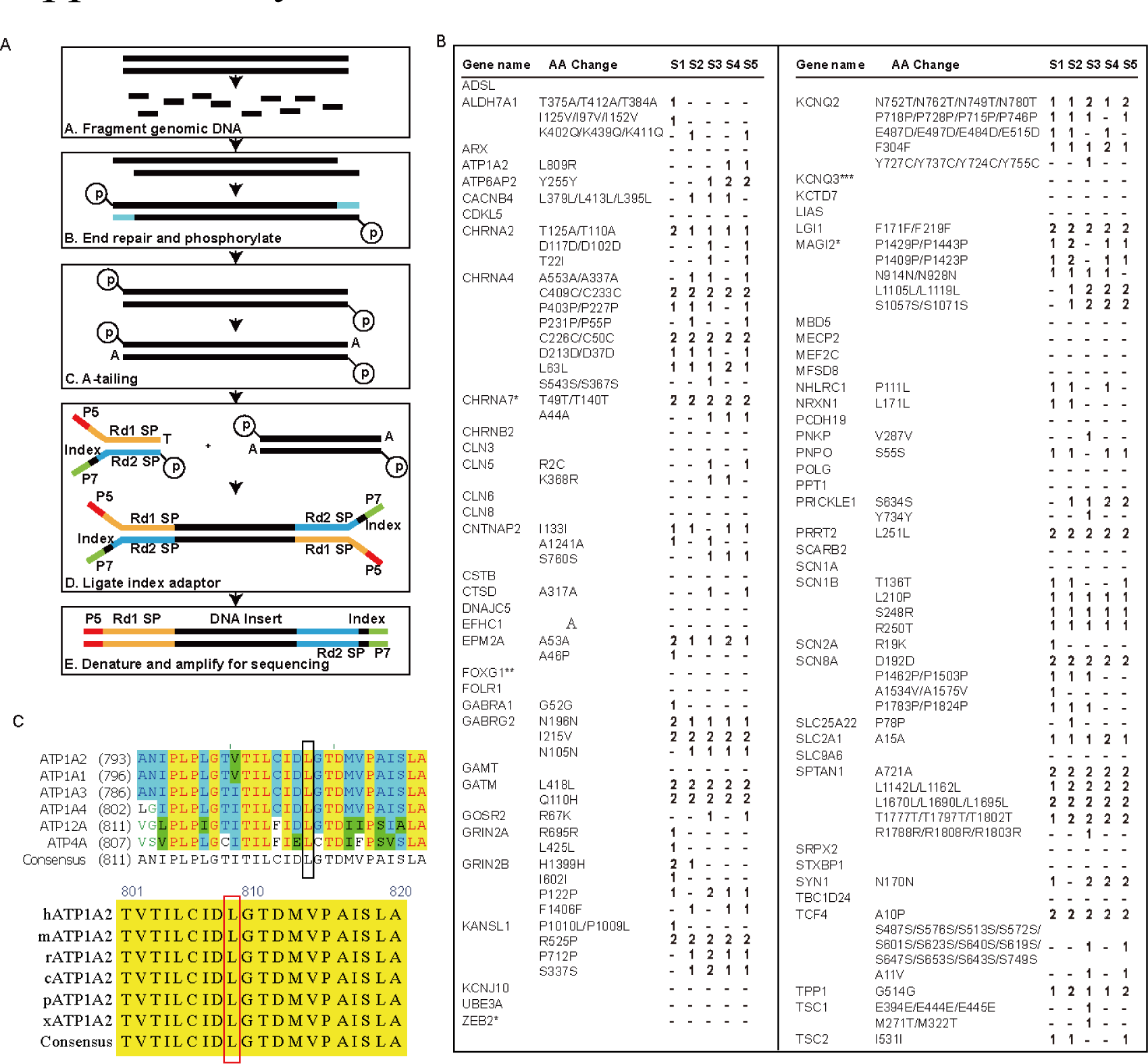
Strategy of the whole-exome sequencing and conserved p.L809 among species. **A,**Schematic showing the whole-exome sequencing. **B**, Epilepsy gene analysis to screen the susceptibility genes for epilepsy. 1, heterozygote; 2, homozygote; -, normal; S1, I-1; S2, II-2; S3, II-3; S4, III-4; S5, III-5. **C**, The ClustalX comparison of amino acid sequences for the K^+^-binding S6 segment shows full conservation of leu809 (arrow) across family proteins (upper) and different species (bottom). This leucine 809 residue (highlighted with a rectangle) is highly conservedwithin families and across species. h, human; m, mouse; r, rat; c, chicken; p, pig; x, *Xenopus*.

**Supplementary Fig. 2.**
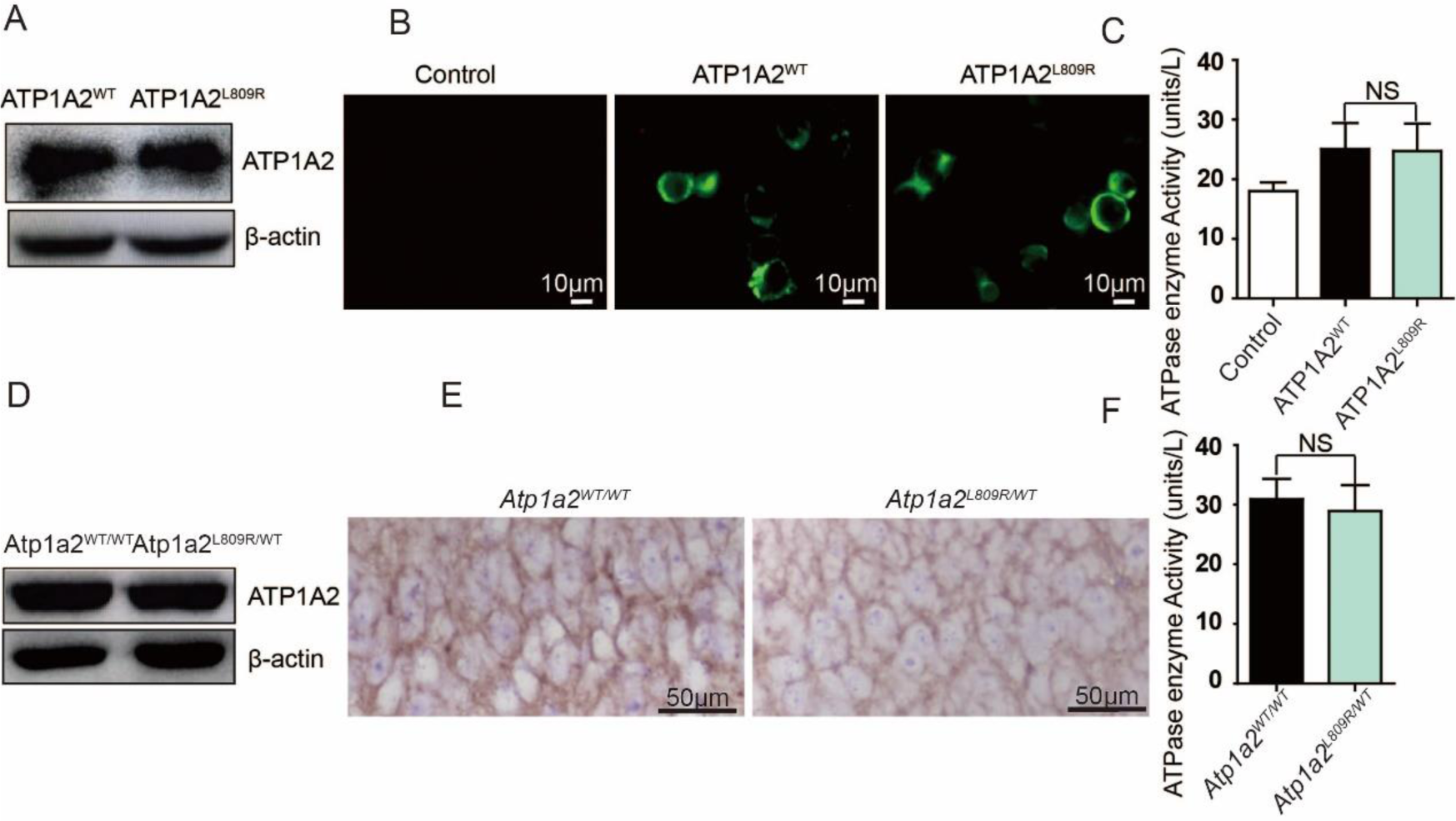
Protein level, membrane localization, and ATPase activity of ATP1A2 were not affected by p.L809R mutation. **A**, Western blot analysis of ATP1A2 in 293T cells transfected with *ATP1A2^WT^* and *ATP1A2^L809R^*. **B**, Immunofluorescent images revealed the membrane distribution of ATP1A2^WT^ and ATP1A2^L809R^ in 293T cells transfected with pEGFP-C1-ATP1A2^WT^ or pEGFP-C1-ATP1A2^L809R^. Scale bar, 10 μm. **C**, ATPase activity analysis of ATP1A2 in 293T cells showed no significant (NS) difference between cells transfected with *ATP1A2^WT^* and *ATP1A2^L809R^*. Data are presented as mean ± SD. No statistical significance by one-way ANOVA with Dunett’s post hoc test. **D**, Western blot analysis of ATP1A2 in the brain tissue of *Atp1a2^WT/WT^* and *Atp1a2^L809R/WT^* mice. **E**, Immunohistochemistry images showing membrane distribution of ATP1A2 in the brain tissue of *Atp1a2^WT/WT^* and *Atp1a2^L809R/WT^* mice. Scale bar, 50 μm. **F**, ATPase activity of ATP1A2 in the brain tissue showed no significant (NS) difference between *Atp1a2^WT/WT^* and *Atp1a2^L809R/WT^* mice. n=6. Data are presented as mean ± SD. No statistical significance by unpaired t-test.

**Supplementary Fig. 3.**
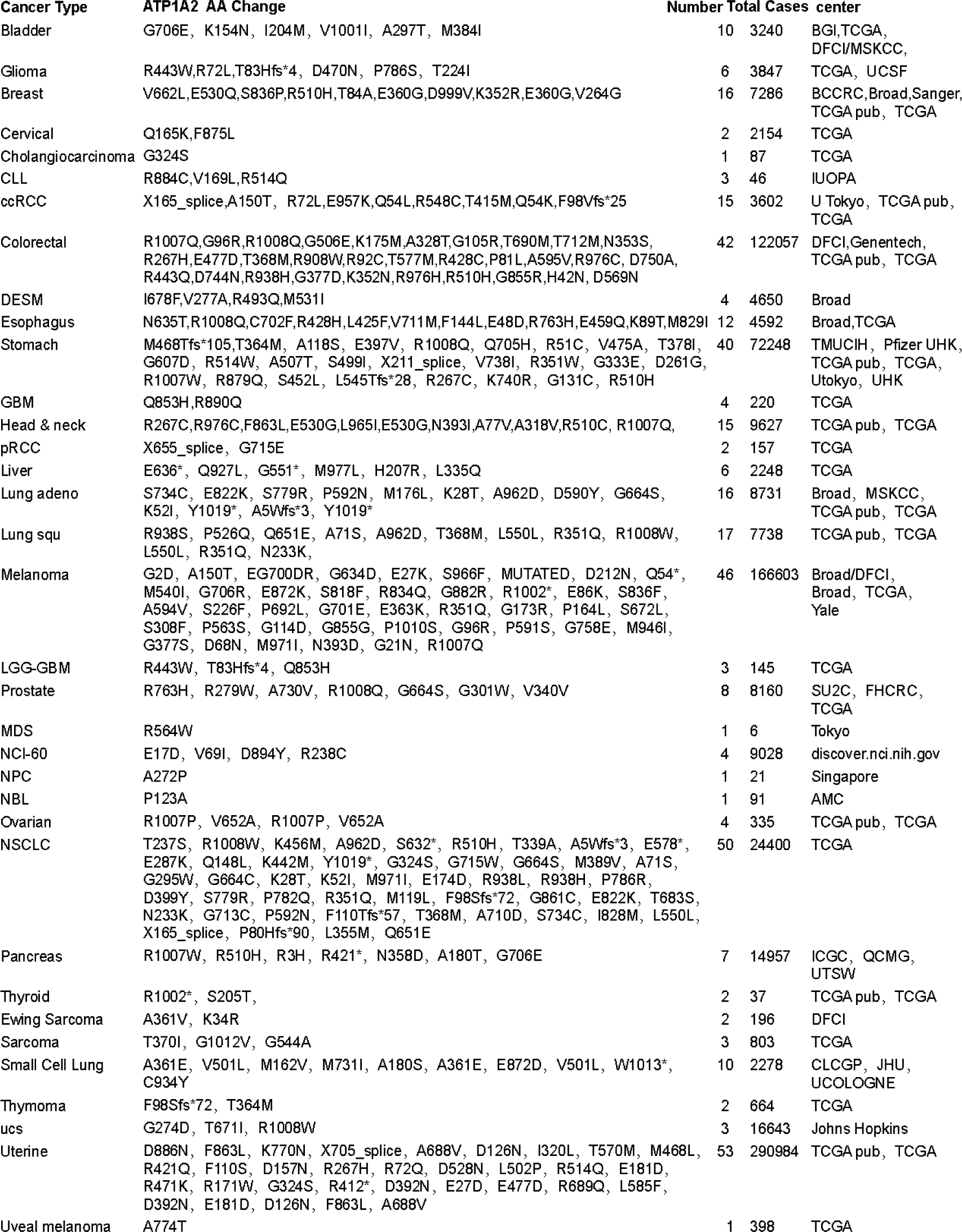
Diseases caused by *ATP1A2* mutation. Frequent mutations in *ATP1A2* genes uncovered from 35 different types of human cancers. Information about mutations in the *ATP1A2* gene was extracted from TCGA (The Cancer Genome Atlas) datasets(Gao et al., 2013) (Cerami et al., 2012). Mutations in the *ATP1A2* gene are indicated as amino acid alteration or distinct mutation types: Nonsense (*), Missense, or Splice. The center reports the mutation, number of mutations found, and number of cases for each cancer type sequenced. CLL, chronic lymphocytic leukemia; ccRCC, clear cell renal cell carcinoma; DESM, desmoplastic melanoma; GBM, glioblastoma multiforme; pRCC, papillary renal cell carcinoma; LGG-GBM, low-grade glioma-glioblastoma multiforme; MDS, myelodysplastic syndrome; NCI-60, national cancer institute’s collection of 60 human cancerous cell lines; NPC, neuroendocrine prostate cancer; NBL, neuroblastoma; NSCLC, non-small-cell lung carcinoma; UCS, uterine carcinosarcoma.

**Supplementary Video 1 Representative video of the ictal behavior of *Atp1a2^L809R/WT^* mice.**

